## Supplementary File 2 for "Community diversity is associated with intra-species genetic diversity and gene loss in the human gut microbiome"

GLMMs and GAMs summary

Naima Madi

19/10/2022

### 1. GAMs with polymorphism rate at synonymous sites (S) in a focal species as a function of community diversity

#### 1.1. Polymorphism rate (S) as a function of shannon diversity

```
##
## Family: Beta regression(83.127)
## Link function: logit
##
## Formula:
## polymorphism_rate ~ s(species_name, Shannon_alpha_diversity,
##     bs = "fs") + s(sample_id, bs = "re") + s(subject_id, bs = "re") +
##     s(Shannon_alpha_diversity) + s(total_reads_orig)
##
## Parametric coefficients:
##             Estimate Std. Error z value Pr(>|z|)
## (Intercept) -5.50462    0.06222  -88.47   <2e-16 ***
## ---
## Signif. codes:  0 '***' 0.001 '**' 0.01 '*' 0.05 '.' 0.1 ' ' 1
##
## Approximate significance of smooth terms:
##                                              edf  Ref.df   Chi.sq  p-value
## s(species_name,Shannon_alpha_diversity)  94.2839 657.000 3045.219  < 2e-16 ***
## s(sample_id)                              0.1071 464.000    0.064 1.000000
## s(subject_id)                           126.8345 248.000  341.220  < 2e-16 ***
## s(Shannon_alpha_diversity)                1.0654   1.110    4.687 0.031412 *
## s(total_reads_orig)                       2.1916   2.622   15.605 0.000883 ***
## ---
## Signif. codes:  0 '***' 0.001 '**' 0.01 '*' 0.05 '.' 0.1 ' ' 1
##
## R-sq.(adj) =  0.198   Deviance explained = 33.2%
## -REML = -44506  Scale est. = 1         n = 8324
```

#### 1.2. Polymorphism rate (S) as a function of species richness (estimated on all data)

```
##
## Family: Beta regression(82.337)
## Link function: logit
##
## Formula:
## polymorphism_rate ~ s(species_richness) + s(total_reads_orig) +
##     s(species_name, species_richness, bs = "fs") + s(sample_id,
##     bs = "re") + s(subject_id, bs = "re")
##
## Parametric coefficients:
##             Estimate Std. Error z value Pr(>|z|)
## (Intercept) -5.50343    0.06384  -86.21   <2e-16 ***
## ---
## Signif. codes:  0 '***' 0.001 '**' 0.01 '*' 0.05 '.' 0.1 ' ' 1
##
## Approximate significance of smooth terms:
##                                       edf  Ref.df   Chi.sq  p-value
## s(species_richness)                2.3921   2.897   10.596   0.0172 *
## s(total_reads_orig)                2.6499   3.180   29.839 2.32e-06 ***
## s(species_name,species_richness)  74.3303 653.000 2899.471  < 2e-16 ***
## s(sample_id)                       0.1199 464.000    0.069   1.0000
## s(subject_id)                    129.6311 248.000  355.976  < 2e-16 ***
## ---
## Signif. codes:  0 '***' 0.001 '**' 0.01 '*' 0.05 '.' 0.1 ' ' 1
##
## R-sq.(adj) =  0.191   Deviance explained = 32.3%
## -REML = -44490  Scale est. = 1         n = 8324
```

#### 1.3. Polymorphism rate (S) as a function of species richness (estimated on rarefied data)

```
##
## Family: Beta regression(81.16)
## Link function: logit
##
## Formula:
## polymorphism_rate ~ s(species_richness) + s(total_reads_orig) +
##     s(species_name, species_richness, bs = "fs") + s(sample_id,
##     bs = "re") + s(subject_id, bs = "re")
##
## Parametric coefficients:
##             Estimate Std. Error z value Pr(>|z|)
## (Intercept) -5.49604    0.06379  -86.16   <2e-16 ***
## ---
## Signif. codes:  0 '***' 0.001 '**' 0.01 '*' 0.05 '.' 0.1 ' ' 1
##
## Approximate significance of smooth terms:
##                                        edf  Ref.df   Chi.sq  p-value
## s(species_richness)                2.68466   3.292   20.199 0.000263 ***
## s(total_reads_orig)                1.97641   2.344    8.043 0.020662 *
## s(species_name,species_richness)  64.97097 664.000 2823.442  < 2e-16 ***
## s(sample_id)                       0.03536 434.000    0.021 1.000000
## s(subject_id)                    121.68267 234.000  319.502  < 2e-16 ***
## ---
## Signif. codes:  0 '***' 0.001 '**' 0.01 '*' 0.05 '.' 0.1 ' ' 1
##
## R-sq.(adj) =   0.19   Deviance explained = 32.2%
## -REML = -43612  Scale est. = 1         n = 8153
```

### 2. GAMs with polymorphism rate at synonymous sites in a focal species as a function of shannon diversity at higher taxonomic levels

#### 2.1. Polymorphism (S) rate as a function of phylum shannon diversity

```
##
## Family: Beta regression(82.102)
## Link function: logit
##
## Formula:
## polymorphism_rate ~ s(species_name, phyla_shannon, bs = "fs") +
##     s(sample_id, bs = "re") + s(subject_id, bs = "re") + s(phyla_shannon) +
##     s(total_reads_orig)
##
## Parametric coefficients:
##             Estimate Std. Error z value Pr(>|z|)
## (Intercept) -5.50443    0.06413  -85.83   <2e-16 ***
## ---
## Signif. codes:  0 '***' 0.001 '**' 0.01 '*' 0.05 '.' 0.1 ' ' 1
##
## Approximate significance of smooth terms:
##                                    edf  Ref.df   Chi.sq  p-value
## s(species_name,phyla_shannon)  63.0641 667.000 2845.917  < 2e-16 ***
## s(sample_id)                    0.0525 464.000    0.030 1.000000
## s(subject_id)                 135.0904 248.000  391.928  < 2e-16 ***
## s(phyla_shannon)                1.0008   1.001    0.399 0.528475
## s(total_reads_orig)             2.2930   2.750   20.019 0.000224 ***
## ---
## Signif. codes:  0 '***' 0.001 '**' 0.01 '*' 0.05 '.' 0.1 ' ' 1
##
## R-sq.(adj) =  0.189   Deviance explained =   32%
## -REML = -44485  Scale est. = 1         n = 8324
```

#### 2.2. Polymorphism rate (S) as a function of class shannon diversity

```
##
## Family: Beta regression(82.102)
## Link function: logit
##
## Formula:
## polymorphism_rate ~ s(species_name, class_shannon, bs = "fs") +
##     s(sample_id, bs = "re") + s(subject_id, bs = "re") + s(class_shannon) +
##     s(total_reads_orig)
##
## Parametric coefficients:
##             Estimate Std. Error z value Pr(>|z|)
## (Intercept) -5.50443    0.06413  -85.83   <2e-16 ***
## ---
## Signif. codes:  0 '***' 0.001 '**' 0.01 '*' 0.05 '.' 0.1 ' ' 1
##
## Approximate significance of smooth terms:
##                                     edf  Ref.df   Chi.sq  p-value
## s(species_name,class_shannon)  63.07154 668.000 2846.265  < 2e-16 ***
## s(sample_id)                    0.05227 464.000    0.030 1.000000
## s(subject_id)                 135.07734 248.000  391.858  < 2e-16 ***
## s(class_shannon)                1.00078   1.001    0.418 0.518425
## s(total_reads_orig)             2.29239   2.749   19.996 0.000227 ***
## ---
## Signif. codes:  0 '***' 0.001 '**' 0.01 '*' 0.05 '.' 0.1 ' ' 1
##
## R-sq.(adj) =  0.189   Deviance explained =   32%
## -REML = -44485  Scale est. = 1         n = 8324
```

#### 2.3. Polymorphism rate (S) as a function of order shannon diversity

```
##
## Family: Beta regression(82.095)
## Link function: logit
##
## Formula:
## polymorphism_rate ~ s(species_name, order_shannon, bs = "fs") +
##     s(sample_id, bs = "re") + s(subject_id, bs = "re") + s(order_shannon) +
##     s(total_reads_orig)
##
## Parametric coefficients:
##             Estimate Std. Error z value Pr(>|z|)
## (Intercept) -5.50422    0.06409  -85.88   <2e-16 ***
## ---
## Signif. codes:  0 '***' 0.001 '**' 0.01 '*' 0.05 '.' 0.1 ' ' 1
##
## Approximate significance of smooth terms:
##                                     edf  Ref.df   Chi.sq  p-value
## s(species_name,order_shannon)  63.05687 667.000 2836.488  < 2e-16 ***
## s(sample_id)                    0.03421 464.000    0.019 1.000000
## s(subject_id)                 134.82575 248.000  389.563  < 2e-16 ***
## s(order_shannon)                1.00290   1.005    0.732 0.394604
## s(total_reads_orig)             2.28820   2.744   19.891 0.000235 ***
## ---
## Signif. codes:  0 '***' 0.001 '**' 0.01 '*' 0.05 '.' 0.1 ' ' 1
##
## R-sq.(adj) =  0.189   Deviance explained =   32%
## -REML = -44485  Scale est. = 1         n = 8324
```

#### 2.4. Polymorphism rate (S) as a function of family shannon diversity

```
##
## Family: Beta regression(82.6)
## Link function: logit
##
## Formula:
## polymorphism_rate ~ s(species_name, family_shannon, bs = "fs") +
##     s(sample_id, bs = "re") + s(subject_id, bs = "re") + s(family_shannon) +
##     s(total_reads_orig)
##
## Parametric coefficients:
##             Estimate Std. Error z value Pr(>|z|)
## (Intercept) -5.50114    0.06373  -86.32   <2e-16 ***
## ---
## Signif. codes:  0 '***' 0.001 '**' 0.01 '*' 0.05 '.' 0.1 ' ' 1
##
## Approximate significance of smooth terms:
##                                     edf  Ref.df   Chi.sq  p-value
## s(species_name,family_shannon)  81.6475 665.000 2929.823  < 2e-16 ***
## s(sample_id)                     0.0644 464.000    0.038 1.000000
## s(subject_id)                  128.9861 248.000  351.108  < 2e-16 ***
## s(family_shannon)                1.0027   1.005    4.676 0.030850 *
## s(total_reads_orig)              2.1864   2.617   15.737 0.000807 ***
## ---
## Signif. codes:  0 '***' 0.001 '**' 0.01 '*' 0.05 '.' 0.1 ' ' 1
##
## R-sq.(adj) =  0.192   Deviance explained = 32.6%
## -REML = -44497  Scale est. = 1         n = 8324
```

#### 2.5. Polymorphism rate (S) as a function of genus shannon diversity

```
##
## Family: Beta regression(82.709)
## Link function: logit
##
## Formula:
## polymorphism_rate ~ s(species_name, genus_shannon, bs = "fs") +
##     s(sample_id, bs = "re") + s(subject_id, bs = "re") + s(genus_shannon) +
##     s(total_reads_orig)
##
## Parametric coefficients:
##             Estimate Std. Error z value Pr(>|z|)
## (Intercept)  -5.5000     0.0636  -86.48   <2e-16 ***
## ---
## Signif. codes:  0 '***' 0.001 '**' 0.01 '*' 0.05 '.' 0.1 ' ' 1
##
## Approximate significance of smooth terms:
##                                     edf  Ref.df   Chi.sq p-value
## s(species_name,genus_shannon)  84.64871 665.000 2937.879 < 2e-16 ***
## s(sample_id)                    0.04288 464.000    0.026 1.00000
## s(subject_id)                 125.68401 248.000  330.285 < 2e-16 ***
## s(genus_shannon)                1.00319   1.006    6.506 0.01093 *
## s(total_reads_orig)             2.17477   2.603   15.552 0.00086 ***
## ---
## Signif. codes:  0 '***' 0.001 '**' 0.01 '*' 0.05 '.' 0.1 ' ' 1
##
## R-sq.(adj) =  0.193   Deviance explained = 32.7%
## -REML = -44502  Scale est. = 1         n = 8324
```

### 3. GAMs with polymorphism rate (at synonymous sites) in a focal species as a function of richness at higher taxonomic levels

#### 3.1. Polymorphism rate (S) as a function of phylum richness

```
##
## Family: Beta regression(82.85)
## Link function: logit
##
## Formula:
## polymorphism_rate ~ s(species_name, phyla_nb, bs = "fs", k = 5) +
##     s(sample_id, bs = "re", k = 5) + s(subject_id, bs = "re",
##     k = 5) + s(phyla_nb, k = 5) + s(total_reads_orig, k = 5)
##
## Parametric coefficients:
##             Estimate Std. Error z value Pr(>|z|)
## (Intercept) -5.50599    0.06421  -85.75   <2e-16 ***
## ---
## Signif. codes:  0 '***' 0.001 '**' 0.01 '*' 0.05 '.' 0.1 ' ' 1
##
## Approximate significance of smooth terms:
##                                edf  Ref.df   Chi.sq  p-value
## s(species_name,phyla_nb)  84.18112 332.000 3039.620  < 2e-16 ***
## s(sample_id)               0.05118 464.000    0.029  1.00000
## s(subject_id)            129.94503 248.000  356.246  < 2e-16 ***
## s(phyla_nb)                1.00193   1.003    7.517  0.00614 **
## s(total_reads_orig)        2.19187   2.550   24.352 2.07e-05 ***
## ---
## Signif. codes:  0 '***' 0.001 '**' 0.01 '*' 0.05 '.' 0.1 ' ' 1
##
## R-sq.(adj) =  0.196   Deviance explained = 32.8%
## -REML = -44497  Scale est. = 1         n = 8324
```

#### 3.2. Polymorphism rate (S) as a function of class richness

```
##
## Family: Beta regression(82.57)
## Link function: logit
##
## Formula:
## polymorphism_rate ~ s(species_name, class_nb, bs = "fs", k = 5) +
##     s(sample_id, bs = "re", k = 5) + s(subject_id, bs = "re",
##     k = 5) + s(class_nb, k = 5) + s(total_reads_orig, k = 5)
##
## Parametric coefficients:
##             Estimate Std. Error z value Pr(>|z|)
## (Intercept) -5.50526    0.06414  -85.83   <2e-16 ***
## ---
## Signif. codes:  0 '***' 0.001 '**' 0.01 '*' 0.05 '.' 0.1 ' ' 1
##
## Approximate significance of smooth terms:
##                                edf  Ref.df   Chi.sq  p-value
## s(species_name,class_nb) 7.888e+01 336.000 2985.657  < 2e-16 ***
## s(sample_id)             3.942e-03 464.000    0.002   1.0000
## s(subject_id)            1.317e+02 248.000  368.973  < 2e-16 ***
## s(class_nb)              1.003e+00   1.005    6.501   0.0109 *
## s(total_reads_orig)      2.254e+00   2.618   26.160 1.23e-05 ***
## ---
## Signif. codes:  0 '***' 0.001 '**' 0.01 '*' 0.05 '.' 0.1 ' ' 1
##
## R-sq.(adj) =  0.194   Deviance explained = 32.5%
## -REML = -44492  Scale est. = 1         n = 8324
```

#### 3.3. Polymorphism rate (S) as a function of order richness

```
##
## Family: Beta regression(82.285)
## Link function: logit
##
## Formula:
## polymorphism_rate ~ s(species_name, order_nb, bs = "fs", k = 5) +
##     s(sample_id, bs = "re", k = 5) + s(subject_id, bs = "re",
##     k = 5) + s(order_nb, k = 5) + s(total_reads_orig, k = 5)
##
## Parametric coefficients:
##             Estimate Std. Error z value Pr(>|z|)
## (Intercept) -5.50518    0.06406  -85.94   <2e-16 ***
## ---
## Signif. codes:  0 '***' 0.001 '**' 0.01 '*' 0.05 '.' 0.1 ' ' 1
##
## Approximate significance of smooth terms:
##                              edf  Ref.df   Chi.sq  p-value
## s(species_name,order_nb)  72.524 337.000 2913.982  < 2e-16 ***
## s(sample_id)               0.039 464.000    0.022   1.0000
## s(subject_id)            132.918 248.000  372.550  < 2e-16 ***
## s(order_nb)                1.009   1.015    5.628   0.0181 *
## s(total_reads_orig)        2.214   2.576   25.006 1.01e-05 ***
## ---
## Signif. codes:  0 '***' 0.001 '**' 0.01 '*' 0.05 '.' 0.1 ' ' 1
##
## R-sq.(adj) =   0.19   Deviance explained = 32.2%
## -REML = -44488  Scale est. = 1         n = 8324
```

#### 3.4. Polymorphism rate (S) as a function of family richness

```
##
## Family: Beta regression(82.251)
## Link function: logit
##
## Formula:
## polymorphism_rate ~ s(species_name, family_nb, bs = "fs", k = 5) +
##     s(sample_id, bs = "re", k = 5) + s(subject_id, bs = "re",
##     k = 5) + s(family_nb, k = 5) + s(total_reads_orig, k = 5)
##
## Parametric coefficients:
##             Estimate Std. Error z value Pr(>|z|)
## (Intercept) -5.50380    0.06389  -86.15   <2e-16 ***
## ---
## Signif. codes:  0 '***' 0.001 '**' 0.01 '*' 0.05 '.' 0.1 ' ' 1
##
## Approximate significance of smooth terms:
##                                 edf  Ref.df   Chi.sq  p-value
## s(species_name,family_nb)  71.93665 338.000 2902.469  < 2e-16 ***
## s(sample_id)                0.03462 464.000    0.020  1.00000
## s(subject_id)             130.42838 248.000  354.350  < 2e-16 ***
## s(family_nb)                1.00032   1.001    9.753  0.00179 **
## s(total_reads_orig)         2.24867   2.615   27.497 3.37e-06 ***
## ---
## Signif. codes:  0 '***' 0.001 '**' 0.01 '*' 0.05 '.' 0.1 ' ' 1
##
## R-sq.(adj) =  0.191   Deviance explained = 32.2%
## -REML = -44490  Scale est. = 1         n = 8324
```

#### 3.5. Polymorphism rate (S) as a function of genus richness

```
##
## Family: Beta regression(82.453)
## Link function: logit
##
## Formula:
## polymorphism_rate ~ s(species_name, genus_nb, bs = "fs", k = 5) +
##     s(sample_id, bs = "re", k = 5) + s(subject_id, bs = "re",
##     k = 5) + s(genus_nb, k = 5) + s(total_reads_orig, k = 5)
##
## Parametric coefficients:
##             Estimate Std. Error z value Pr(>|z|)
## (Intercept) -5.50417    0.06381  -86.26   <2e-16 ***
## ---
## Signif. codes:  0 '***' 0.001 '**' 0.01 '*' 0.05 '.' 0.1 ' ' 1
##
## Approximate significance of smooth terms:
##                                edf  Ref.df   Chi.sq  p-value
## s(species_name,genus_nb)  78.66673 338.000 2923.286  < 2e-16 ***
## s(sample_id)               0.01746 464.000    0.010  1.00000
## s(subject_id)            129.82405 248.000  353.898  < 2e-16 ***
## s(genus_nb)                1.00718   1.012    8.095  0.00448 **
## s(total_reads_orig)        2.28861   2.659   28.260 2.42e-06 ***
## ---
## Signif. codes:  0 '***' 0.001 '**' 0.01 '*' 0.05 '.' 0.1 ' ' 1
##
## R-sq.(adj) =  0.193   Deviance explained = 32.4%
## -REML = -44492  Scale est. = 1         n = 8324
```

### 4. GAMs with polymorphism rate at non synonymous sites (NS) in a focal species as a function of community diversity

#### 4.1. Polymorphism rate (NS) as a function of shannon diversity

```
##
## Family: Beta regression(854.674)
## Link function: logit
##
## Formula:
## polymorphism_nonsyn ~ s(species, alpha_div, bs = "fs") + s(sample_id,
##     bs = "re") + s(subject, bs = "re") + s(alpha_div) + s(total_reads_orig)
##
## Parametric coefficients:
##             Estimate Std. Error z value Pr(>|z|)
## (Intercept) -7.40965    0.05846  -126.8   <2e-16 ***
## ---
## Signif. codes:  0 '***' 0.001 '**' 0.01 '*' 0.05 '.' 0.1 ' ' 1
##
## Approximate significance of smooth terms:
##                           edf  Ref.df   Chi.sq p-value
## s(species,alpha_div) 121.6380 668.000 3426.332  <2e-16 ***
## s(sample_id)           0.3785 464.000    0.313   0.989
## s(subject)           145.1280 248.000  468.255  <2e-16 ***
## s(alpha_div)           1.0336   1.056    0.070   0.842
## s(total_reads_orig)    3.2545   3.923   92.304  <2e-16 ***
## ---
## Signif. codes:  0 '***' 0.001 '**' 0.01 '*' 0.05 '.' 0.1 ' ' 1
##
## R-sq.(adj) =  0.248   Deviance explained = 33.6%
## -REML = -56566  Scale est. = 1         n = 8324
```

#### 4.2. Polymorphism rate (NS) as a function of species richness (all data)

```
##
## Family: Beta regression(848.533)
## Link function: logit
##
## Formula:
## polymorphism_nonsyn ~ s(richness) + s(total_reads_orig) + s(species,
##     richness, bs = "fs") + s(sample_id, bs = "re") + s(subject,
##     bs = "re")
##
## Parametric coefficients:
##             Estimate Std. Error z value Pr(>|z|)
## (Intercept) -7.40651    0.05218  -141.9   <2e-16 ***
## ---
## Signif. codes:  0 '***' 0.001 '**' 0.01 '*' 0.05 '.' 0.1 ' ' 1
##
## Approximate significance of smooth terms:
##                          edf  Ref.df   Chi.sq p-value
## s(richness)           3.5898   4.301    8.238  0.0928 .
## s(total_reads_orig)   3.6766   4.421  109.187  <2e-16 ***
## s(species,richness) 123.6962 667.000 3233.935  <2e-16 ***
## s(sample_id)          0.1284 464.000    0.103  0.9963
## s(subject)          140.2825 248.000  432.162  <2e-16 ***
## ---
## Signif. codes:  0 '***' 0.001 '**' 0.01 '*' 0.05 '.' 0.1 ' ' 1
##
## R-sq.(adj) =  0.243   Deviance explained = 33.1%
## -REML = -56534  Scale est. = 1         n = 8324
```

#### 4.3. Polymorphism rate (NS) as a function of species richness (rarefied data)

```
##
## Family: Beta regression(851.252)
## Link function: logit
##
## Formula:
## polymorphism_nonsyn ~ s(richness_rare) + s(total_reads_orig) +
##     s(species, richness_rare, bs = "fs") + s(sample_id, bs = "re") +
##     s(subject, bs = "re")
##
## Parametric coefficients:
##             Estimate Std. Error z value Pr(>|z|)
## (Intercept) -7.40824    0.06345  -116.8   <2e-16 ***
## ---
## Signif. codes:  0 '***' 0.001 '**' 0.01 '*' 0.05 '.' 0.1 ' ' 1
##
## Approximate significance of smooth terms:
##                               edf  Ref.df   Chi.sq p-value
## s(richness_rare)           1.5383   1.807    1.985   0.257
## s(total_reads_orig)        3.4215   4.124   92.685  <2e-16 ***
## s(species,richness_rare) 117.7944 665.000 3371.510  <2e-16 ***
## s(sample_id)               0.2617 464.000    0.220   0.981
## s(subject)               143.7846 248.000  445.570  <2e-16 ***
## ---
## Signif. codes:  0 '***' 0.001 '**' 0.01 '*' 0.05 '.' 0.1 ' ' 1
##
## R-sq.(adj) =  0.244   Deviance explained = 33.3%
## -REML = -56567  Scale est. = 1         n = 8324
```

### 5. GAMs with polymorphism rate (at non synonymous sites) in a focal species as a function of shannon diversity at higher taxonomic levels

#### 5.1. Polymorphism rate (NS) as a function of phylum shannon diversity

```
##
## Family: Beta regression(850.356)
## Link function: logit
##
## Formula:
## polymorphism_nonsyn ~ s(species, phyla_shannon, bs = "fs") +
##     s(sample_id, bs = "re") + s(subject, bs = "re") + s(phyla_shannon) +
##     s(total_reads_orig)
##
## Parametric coefficients:
##             Estimate Std. Error z value Pr(>|z|)
## (Intercept) -7.41389    0.06201  -119.6   <2e-16 ***
## ---
## Signif. codes:  0 '***' 0.001 '**' 0.01 '*' 0.05 '.' 0.1 ' ' 1
##
## Approximate significance of smooth terms:
##                              edf  Ref.df   Chi.sq p-value
## s(species,phyla_shannon) 103.872 668.000 3269.572  <2e-16 ***
## s(sample_id)               0.135 464.000    0.109   0.995
## s(subject)               150.384 248.000  511.372  <2e-16 ***
## s(phyla_shannon)           1.033   1.058    1.141   0.284
## s(total_reads_orig)        3.310   3.997   97.817  <2e-16 ***
## ---
## Signif. codes:  0 '***' 0.001 '**' 0.01 '*' 0.05 '.' 0.1 ' ' 1
##
## R-sq.(adj) =  0.241   Deviance explained = 33.1%
## -REML = -56564  Scale est. = 1         n = 8324
```

#### 5.2. Polymorphism rate (NS) as a function of class shannon diversity

```
##
## Family: Beta regression(851.796)
## Link function: logit
##
## Formula:
## polymorphism_nonsyn ~ s(species, class_shannon, bs = "fs") +
##     s(sample_id, bs = "re") + s(subject, bs = "re") + s(class_shannon) +
##     s(total_reads_orig)
##
## Parametric coefficients:
##             Estimate Std. Error z value Pr(>|z|)
## (Intercept) -7.40564    0.06189  -119.7   <2e-16 ***
## ---
## Signif. codes:  0 '***' 0.001 '**' 0.01 '*' 0.05 '.' 0.1 ' ' 1
##
## Approximate significance of smooth terms:
##                                edf  Ref.df   Chi.sq p-value
## s(species,class_shannon) 109.85392 668.000 3280.865  <2e-16 ***
## s(sample_id)               0.08542 464.000    0.069   0.995
## s(subject)               151.01909 248.000  514.122  <2e-16 ***
## s(class_shannon)           2.00624   2.457    2.804   0.286
## s(total_reads_orig)        3.28034   3.960  103.538  <2e-16 ***
## ---
## Signif. codes:  0 '***' 0.001 '**' 0.01 '*' 0.05 '.' 0.1 ' ' 1
##
## R-sq.(adj) =   0.24   Deviance explained = 33.4%
## -REML = -56558  Scale est. = 1         n = 8324
```

#### 5.3. Polymorphism rate (NS) as a function of order shannon diversity

```
##
## Family: Beta regression(852.236)
## Link function: logit
##
## Formula:
## polymorphism_nonsyn ~ s(species, order_shannon, bs = "fs") +
##     s(sample_id, bs = "re") + s(subject, bs = "re") + s(order_shannon) +
##     s(total_reads_orig)
##
## Parametric coefficients:
##             Estimate Std. Error z value Pr(>|z|)
## (Intercept) -7.41338    0.06557  -113.1   <2e-16 ***
## ---
## Signif. codes:  0 '***' 0.001 '**' 0.01 '*' 0.05 '.' 0.1 ' ' 1
##
## Approximate significance of smooth terms:
##                               edf  Ref.df   Chi.sq p-value
## s(species,order_shannon)  91.9823 668.000 3291.619  <2e-16 ***
## s(sample_id)               0.1375 464.000    0.109   0.997
## s(subject)               151.2407 248.000  513.477  <2e-16 ***
## s(order_shannon)           2.4651   2.999    5.446   0.138
## s(total_reads_orig)        3.2664   3.942  102.025  <2e-16 ***
## ---
## Signif. codes:  0 '***' 0.001 '**' 0.01 '*' 0.05 '.' 0.1 ' ' 1
##
## R-sq.(adj) =  0.242   Deviance explained = 33.2%
## -REML = -56582  Scale est. = 1         n = 8324
```

#### 5.4. Polymorphism rate (NS) as a function of family shannon diversity

```
##
## Family: Beta regression(851.323)
## Link function: logit
##
## Formula:
## polymorphism_nonsyn ~ s(species, family_shannon, bs = "fs") +
##     s(sample_id, bs = "re") + s(subject, bs = "re") + s(family_shannon) +
##     s(total_reads_orig)
##
## Parametric coefficients:
##             Estimate Std. Error z value Pr(>|z|)
## (Intercept) -7.41676    0.06363  -116.6   <2e-16 ***
## ---
## Signif. codes:  0 '***' 0.001 '**' 0.01 '*' 0.05 '.' 0.1 ' ' 1
##
## Approximate significance of smooth terms:
##                                edf  Ref.df   Chi.sq p-value
## s(species,family_shannon)  90.1645 665.000 3333.078  <2e-16 ***
## s(sample_id)                0.1478 464.000    0.122   0.988
## s(subject)                146.9384 248.000  477.546  <2e-16 ***
## s(family_shannon)           1.0031   1.006    0.041   0.842
## s(total_reads_orig)         3.2506   3.922   90.611  <2e-16 ***
## ---
## Signif. codes:  0 '***' 0.001 '**' 0.01 '*' 0.05 '.' 0.1 ' ' 1
##
## R-sq.(adj) =  0.244   Deviance explained =   33%
## -REML = -56588  Scale est. = 1         n = 8324
```

#### 5.5. Polymorphism rate (NS) as a function of genus shannon diversity

```
##
## Family: Beta regression(855.56)
## Link function: logit
##
## Formula:
## polymorphism_nonsyn ~ s(species, genus_shannon, bs = "fs") +
##     s(sample_id, bs = "re") + s(subject, bs = "re") + s(genus_shannon) +
##     s(total_reads_orig)
##
## Parametric coefficients:
##             Estimate Std. Error z value Pr(>|z|)
## (Intercept) -7.41443    0.06309  -117.5   <2e-16 ***
## ---
## Signif. codes:  0 '***' 0.001 '**' 0.01 '*' 0.05 '.' 0.1 ' ' 1
##
## Approximate significance of smooth terms:
##                               edf  Ref.df   Chi.sq p-value
## s(species,genus_shannon) 105.3751 668.000 3402.444  <2e-16 ***
## s(sample_id)               0.1687 464.000    0.143   0.977
## s(subject)               145.6475 248.000  466.025  <2e-16 ***
## s(genus_shannon)           1.0056   1.009    0.028   0.871
## s(total_reads_orig)        3.2527   3.924   91.715  <2e-16 ***
## ---
## Signif. codes:  0 '***' 0.001 '**' 0.01 '*' 0.05 '.' 0.1 ' ' 1
##
## R-sq.(adj) =  0.247   Deviance explained = 33.5%
## -REML = -56592  Scale est. = 1         n = 8324
```

### 6. GAMs with polymorphism rate (at non synonymous sites) in a focal species as a function of richness at higher taxonomic levels

#### 6.1. Polymorphism rate (NS) as a function of phylum richness

```
##
## Family: Beta regression(849.927)
## Link function: logit
##
## Formula:
## polymorphism_nonsyn ~ s(species, phyla_nb, k = 5, bs = "fs") +
##     s(sample_id, k = 5, bs = "re") + s(subject, k = 5, bs = "re") +
##     s(phyla_nb, k = 5) + s(total_reads_orig, k = 5)
##
## Parametric coefficients:
##             Estimate Std. Error z value Pr(>|z|)
## (Intercept) -7.41984    0.06468  -114.7   <2e-16 ***
## ---
## Signif. codes:  0 '***' 0.001 '**' 0.01 '*' 0.05 '.' 0.1 ' ' 1
##
## Approximate significance of smooth terms:
##                          edf  Ref.df   Chi.sq p-value
## s(species,phyla_nb)  77.4818 332.000 3399.932  <2e-16 ***
## s(sample_id)          0.1337 464.000    0.107  0.9969
## s(subject)          144.3327 248.000  456.184  <2e-16 ***
## s(phyla_nb)           1.0038   1.006   13.105  0.0003 ***
## s(total_reads_orig)   2.9432   3.328  107.353  <2e-16 ***
## ---
## Signif. codes:  0 '***' 0.001 '**' 0.01 '*' 0.05 '.' 0.1 ' ' 1
##
## R-sq.(adj) =  0.246   Deviance explained = 32.7%
## -REML = -56594  Scale est. = 1         n = 8324
```

#### 6.2. Polymorphism rate (NS) as a function of class richness

```
##
## Family: Beta regression(848.574)
## Link function: logit
##
## Formula:
## polymorphism_nonsyn ~ s(species, class_nb, k = 5, bs = "fs") +
##     s(sample_id, bs = "re", k = 5) + s(subject, bs = "re", k = 5) +
##     s(class_nb, k = 5) + s(total_reads_orig, k = 5)
##
## Parametric coefficients:
##             Estimate Std. Error z value Pr(>|z|)
## (Intercept) -7.41616    0.06345  -116.9   <2e-16 ***
## ---
## Signif. codes:  0 '***' 0.001 '**' 0.01 '*' 0.05 '.' 0.1 ' ' 1
##
## Approximate significance of smooth terms:
##                          edf  Ref.df   Chi.sq p-value
## s(species,class_nb)  98.6016 336.000 3365.620  <2e-16 ***
## s(sample_id)          0.1877 464.000    0.149  0.9978
## s(subject)          145.5007 248.000  466.554  <2e-16 ***
## s(class_nb)           1.0027   1.005    5.712  0.0171 *
## s(total_reads_orig)   2.9709   3.353  108.694  <2e-16 ***
## ---
## Signif. codes:  0 '***' 0.001 '**' 0.01 '*' 0.05 '.' 0.1 ' ' 1
##
## R-sq.(adj) =  0.244   Deviance explained = 32.8%
## -REML = -56567  Scale est. = 1         n = 8324
```

#### 6.3. Polymorphism rate (NS) as a function of order richness

```
##
## Family: Beta regression(846.954)
## Link function: logit
##
## Formula:
## polymorphism_nonsyn ~ s(species, order_nb, bs = "fs") + s(sample_id,
##     bs = "re") + s(subject, bs = "re") + s(order_nb) + s(total_reads_orig)
##
## Parametric coefficients:
##             Estimate Std. Error z value Pr(>|z|)
## (Intercept) -7.42185    0.06436  -115.3   <2e-16 ***
## ---
## Signif. codes:  0 '***' 0.001 '**' 0.01 '*' 0.05 '.' 0.1 ' ' 1
##
## Approximate significance of smooth terms:
##                          edf  Ref.df   Chi.sq  p-value
## s(species,order_nb)  68.9809 658.000 3269.251  < 2e-16 ***
## s(sample_id)          0.1326 464.000    0.108 0.994300
## s(subject)          145.7224 248.000  461.058  < 2e-16 ***
## s(order_nb)           1.0352   1.059   11.954 0.000611 ***
## s(total_reads_orig)   3.3771   4.074  108.737  < 2e-16 ***
## ---
## Signif. codes:  0 '***' 0.001 '**' 0.01 '*' 0.05 '.' 0.1 ' ' 1
##
## R-sq.(adj) =  0.244   Deviance explained = 32.4%
## -REML = -56588  Scale est. = 1         n = 8324
```

#### 6.4. Polymorphism rate (NS) as a function of family richness

```
##
## Family: Beta regression(844.979)
## Link function: logit
##
## Formula:
## polymorphism_nonsyn ~ s(species, family_nb, bs = "fs") + s(sample_id,
##     bs = "re") + s(subject, bs = "re") + s(family_nb) + s(total_reads_orig)
##
## Parametric coefficients:
##             Estimate Std. Error z value Pr(>|z|)
## (Intercept) -7.41469    0.05678  -130.6   <2e-16 ***
## ---
## Signif. codes:  0 '***' 0.001 '**' 0.01 '*' 0.05 '.' 0.1 ' ' 1
##
## Approximate significance of smooth terms:
##                           edf  Ref.df   Chi.sq p-value
## s(species,family_nb) 107.4280 664.000 3185.198  <2e-16 ***
## s(sample_id)           0.1441 464.000    0.118   0.991
## s(subject)           140.3559 248.000  419.052  <2e-16 ***
## s(family_nb)           1.5716   1.875    2.211   0.241
## s(total_reads_orig)    3.4335   4.134  108.800  <2e-16 ***
## ---
## Signif. codes:  0 '***' 0.001 '**' 0.01 '*' 0.05 '.' 0.1 ' ' 1
##
## R-sq.(adj) =  0.243   Deviance explained = 32.6%
## -REML = -56551  Scale est. = 1         n = 8324
```

#### 6.5. Polymorphism rate (NS) as a function of genus richness

```
##
## Family: Beta regression(847.977)
## Link function: logit
##
## Formula:
## polymorphism_nonsyn ~ s(species, genus_nb, bs = "fs") + s(sample_id,
##     bs = "re") + s(subject, bs = "re") + s(genus_nb) + s(total_reads_orig)
##
## Parametric coefficients:
##             Estimate Std. Error z value Pr(>|z|)
## (Intercept) -7.40821    0.06158  -120.3   <2e-16 ***
## ---
## Signif. codes:  0 '***' 0.001 '**' 0.01 '*' 0.05 '.' 0.1 ' ' 1
##
## Approximate significance of smooth terms:
##                          edf  Ref.df   Chi.sq p-value
## s(species,genus_nb) 106.7698 661.000 3261.527  <2e-16 ***
## s(sample_id)          0.2152 464.000    0.177   0.991
## s(subject)          143.3437 248.000  444.411  <2e-16 ***
## s(genus_nb)           1.0310   1.052    2.434   0.122
## s(total_reads_orig)   3.4900   4.209  107.829  <2e-16 ***
## ---
## Signif. codes:  0 '***' 0.001 '**' 0.01 '*' 0.05 '.' 0.1 ' ' 1
##
## R-sq.(adj) =  0.243   Deviance explained = 32.9%
## -REML = -56559  Scale est. = 1         n = 8324
```

### 7. GLMMs with strain count in a focal species as a function of community diversity

#### 7.1. Strain count in a focal species as a function of Shannon diversity

```
##  Family: truncated_nbinom2  ( log )
## Formula:
## strain_nb ~ shannon_diversity + total_reads_orig + (1 | species_id)
## Data: datsc
##
##      AIC      BIC   logLik deviance df.resid
##  25218.2  25256.4 -12604.1  25208.2    15350
##
## Random effects:
##
## Conditional model:
##  Groups     Name        Variance Std.Dev.
##  species_id (Intercept) 0.2825   0.5315
## Number of obs: 15355, groups:  species_id, 184
##
## Overdispersion parameter for truncated_nbinom2 family (): 5.71e+05
##
## Conditional model:
##                   Estimate Std. Error z value Pr(>|z|)
## (Intercept)       -0.30024    0.05086  -5.903 3.57e-09 ***
## shannon_diversity  0.05927    0.01178   5.033 4.83e-07 ***
## total_reads_orig   0.03362    0.01087   3.092  0.00199 **
## ---
## Signif. codes:  0 '***' 0.001 '**' 0.01 '*' 0.05 '.' 0.1 ' ' 1
```

#### 7.2. Strain count in a focal species as a function of species richness (all data)

```
##  Family: truncated_nbinom1  ( log )
## Formula:
## strain_nb ~ species_richness + total_reads_orig + (species_richness |
##     species_id) + (1 | sample_id) + (1 | subject_id)
## Data: datsc
##
##      AIC      BIC   logLik deviance df.resid
##  24936.4  25005.1 -12459.2  24918.4    15346
##
## Random effects:
##
## Conditional model:
##  Groups     Name             Variance  Std.Dev. Corr
##  species_id (Intercept)      0.3005843 0.54826
##             species_richness 0.0001944 0.01394  0.76
##  sample_id  (Intercept)      0.0218740 0.14790
##  subject_id (Intercept)      0.0629320 0.25086
## Number of obs: 15355, groups:  species_id, 184; sample_id, 467; subject_id, 249
##
## Overdispersion parameter for truncated_nbinom1 family (): 1.76e-05
##
## Conditional model:
##                  Estimate Std. Error z value Pr(>|z|)
## (Intercept)      -0.32293    0.05560  -5.808 6.31e-09 ***
## species_richness -0.09799    0.02114  -4.636 3.56e-06 ***
## total_reads_orig  0.10792    0.02163   4.990 6.02e-07 ***
## ---
## Signif. codes:  0 '***' 0.001 '**' 0.01 '*' 0.05 '.' 0.1 ' ' 1
```

#### 7.3. Strain count in a focal species as a function of richness (rarefied data)

```
##  Family: truncated_nbinom2  ( log )
## Formula:
## strain_nb ~ species_richness + total_reads_orig + (1 | species_id)
## Data: datsc
##
##      AIC      BIC   logLik deviance df.resid
##  24680.5  24718.5 -12335.2  24670.5    14958
##
## Random effects:
##
## Conditional model:
##  Groups     Name        Variance Std.Dev.
##  species_id (Intercept) 0.2917   0.5401
## Number of obs: 14963, groups:  species_id, 180
##
## Overdispersion parameter for truncated_nbinom2 family (): 2.89e+05
##
## Conditional model:
##                  Estimate Std. Error z value Pr(>|z|)
## (Intercept)      -0.28554    0.05171  -5.522 3.35e-08 ***
## species_richness -0.05129    0.01182  -4.341 1.42e-05 ***
## total_reads_orig  0.01634    0.01118   1.462    0.144
## ---
## Signif. codes:  0 '***' 0.001 '**' 0.01 '*' 0.05 '.' 0.1 ' ' 1
```

### 8. Strain count in a focal species as a function of shannon diversity at higher taxonomic levels

#### 8.1. Strain count in a focal species as a function of phylum shannon diversity

```
##  Family: truncated_nbinom2  ( log )
## Formula:
## strain_nb ~ phyla_shannon + total_reads_orig + (phyla_shannon |
##     species_id) + (1 | sample_id) + (1 | subject_id)
## Data: datsc2
##
##      AIC      BIC   logLik deviance df.resid
##  24949.9  25018.6 -12465.9  24931.9    15346
##
## Random effects:
##
## Conditional model:
##  Groups     Name          Variance Std.Dev. Corr
##  species_id (Intercept)   0.306800 0.55390
##             phyla_shannon 0.001671 0.04087  -0.99
##  sample_id  (Intercept)   0.025953 0.16110
##  subject_id (Intercept)   0.065625 0.25617
## Number of obs: 15355, groups:  species_id, 184; sample_id, 467; subject_id, 249
##
## Overdispersion parameter for truncated_nbinom2 family (): 3.35e+04
##
## Conditional model:
##                  Estimate Std. Error z value Pr(>|z|)
## (Intercept)      -0.32635    0.05658  -5.768 8.01e-09 ***
## phyla_shannon     0.03018    0.01951   1.547  0.12190
## total_reads_orig  0.06520    0.02062   3.162  0.00157 **
## ---
## Signif. codes:  0 '***' 0.001 '**' 0.01 '*' 0.05 '.' 0.1 ' ' 1
```

#### 8.2. Strain count in a focal species as a function of class shannon diversity

```
##  Family: truncated_nbinom2  ( log )
## Formula:
## strain_nb ~ class_shannon + total_reads_orig + (class_shannon |
##     species_id) + (1 | sample_id) + (1 | subject_id)
## Data: datsc2
##
##      AIC      BIC   logLik deviance df.resid
##  24949.7  25018.4 -12465.8  24931.7    15346
##
## Random effects:
##
## Conditional model:
##  Groups     Name          Variance Std.Dev. Corr
##  species_id (Intercept)   0.308816 0.5557
##             class_shannon 0.001823 0.0427   -1.00
##  sample_id  (Intercept)   0.026762 0.1636
##  subject_id (Intercept)   0.065165 0.2553
## Number of obs: 15355, groups:  species_id, 184; sample_id, 467; subject_id, 249
##
## Overdispersion parameter for truncated_nbinom2 family (): 3.56e+04
##
## Conditional model:
##                  Estimate Std. Error z value Pr(>|z|)
## (Intercept)      -0.32521    0.05670  -5.736  9.7e-09 ***
## class_shannon     0.03187    0.01956   1.629  0.10333
## total_reads_orig  0.06615    0.02067   3.201  0.00137 **
## ---
## Signif. codes:  0 '***' 0.001 '**' 0.01 '*' 0.05 '.' 0.1 ' ' 1
```

#### 8.3. Strain count in a focal species as a function of order shannon diversity

```
##  Family: truncated_nbinom2  ( log )
## Formula:
## strain_nb ~ order_shannon + total_reads_orig + (order_shannon |
##     species_id)
## Data: datsc2
##
##      AIC      BIC   logLik deviance df.resid
##  25235.2  25288.7 -12610.6  25221.2    15348
##
## Random effects:
##
## Conditional model:
##  Groups     Name          Variance Std.Dev. Corr
##  species_id (Intercept)   0.292881 0.54118
##             order_shannon 0.002037 0.04513  -1.00
## Number of obs: 15355, groups:  species_id, 184
##
## Overdispersion parameter for truncated_nbinom2 family (): 3.54e+09
##
## Conditional model:
##                  Estimate Std. Error z value Pr(>|z|)
## (Intercept)      -0.29546    0.05197  -5.685 1.31e-08 ***
## order_shannon     0.03425    0.01359   2.521   0.0117 *
## total_reads_orig  0.02811    0.01092   2.575   0.0100 *
## ---
## Signif. codes:  0 '***' 0.001 '**' 0.01 '*' 0.05 '.' 0.1 ' ' 1
```

#### 8.4. Strain count in a focal species as a function of family shannon diversity

```
##  Family: truncated_nbinom2  ( log )
## Formula:
## strain_nb ~ family_shannon + total_reads_orig + (family_shannon |
##     species_id) + (1 | sample_id) + (1 | subject_id)
## Data: datsc2
##
##      AIC      BIC   logLik deviance df.resid
##  24952.5  25021.3 -12467.3  24934.5    15346
##
## Random effects:
##
## Conditional model:
##  Groups     Name           Variance Std.Dev. Corr
##  species_id (Intercept)    0.306320 0.5535
##             family_shannon 0.001289 0.0359   -0.99
##  sample_id  (Intercept)    0.026419 0.1625
##  subject_id (Intercept)    0.064846 0.2546
## Number of obs: 15355, groups:  species_id, 184; sample_id, 467; subject_id, 249
##
## Overdispersion parameter for truncated_nbinom2 family (): 4e+04
##
## Conditional model:
##                  Estimate Std. Error z value Pr(>|z|)
## (Intercept)      -0.32556    0.05635  -5.777 7.59e-09 ***
## family_shannon    0.02981    0.01966   1.516  0.12952
## total_reads_orig  0.06610    0.02062   3.206  0.00135 **
## ---
## Signif. codes:  0 '***' 0.001 '**' 0.01 '*' 0.05 '.' 0.1 ' ' 1
```

#### 8.5. Strain count in a focal species as a function of genus shannon diversity

```
##  Family: truncated_nbinom2  ( log )
## Formula:
## strain_nb ~ genus_shannon + total_reads_orig + (genus_shannon |
##     species_id) + (1 | sample_id) + (1 | subject_id)
## Data: datsc2
##
##      AIC      BIC   logLik deviance df.resid
##  24940.2  25008.9 -12461.1  24922.2    15346
##
## Random effects:
##
## Conditional model:
##  Groups     Name          Variance Std.Dev. Corr
##  species_id (Intercept)   0.311063 0.55773
##             genus_shannon 0.003269 0.05717  -0.99
##  sample_id  (Intercept)   0.026939 0.16413
##  subject_id (Intercept)   0.062524 0.25005
## Number of obs: 15355, groups:  species_id, 184; sample_id, 467; subject_id, 249
##
## Overdispersion parameter for truncated_nbinom2 family (): 1.11e+08
##
## Conditional model:
##                  Estimate Std. Error z value Pr(>|z|)
## (Intercept)      -0.32776    0.05672  -5.778 7.55e-09 ***
## genus_shannon     0.05746    0.02025   2.837  0.00455 **
## total_reads_orig  0.06556    0.02059   3.184  0.00145 **
## ---
## Signif. codes:  0 '***' 0.001 '**' 0.01 '*' 0.05 '.' 0.1 ' ' 1
```

### 9. GLMMs with strain count in a focal species as a function of richness at higher taxonomic levels

#### 9.1. Strain count in a focal species as a function of phylum richness

```
##  Family: truncated_nbinom2  ( log )
## Formula:          strain_nb ~ phyla_nb + total_reads_orig + (1 | species_id)
## Data: datsc2
##
##      AIC      BIC   logLik deviance df.resid
##  25226.1  25264.3 -12608.1  25216.1    15350
##
## Random effects:
##
## Conditional model:
##  Groups     Name        Variance Std.Dev.
##  species_id (Intercept) 0.2863   0.5351
## Number of obs: 15355, groups:  species_id, 184
##
## Overdispersion parameter for truncated_nbinom2 family (): 1.95e+05
##
## Conditional model:
##                  Estimate Std. Error z value Pr(>|z|)
## (Intercept)      -0.29144    0.05109  -5.705 1.17e-08 ***
## phyla_nb         -0.04854    0.01145  -4.240 2.24e-05 ***
## total_reads_orig  0.03596    0.01102   3.262   0.0011 **
## ---
## Signif. codes:  0 '***' 0.001 '**' 0.01 '*' 0.05 '.' 0.1 ' ' 1
```

#### 9.2. Strain count in a focal species as a function of class richness

```
##  Family: truncated_nbinom1  ( log )
## Formula:          strain_nb ~ class_nb + total_reads_orig + (1 | species_id) +
##     (1 | sample_id) + (1 | subject_id)
## Data: datsc2
##
##      AIC      BIC   logLik deviance df.resid
##  24941.6  24995.1 -12463.8  24927.6    15348
##
## Random effects:
##
## Conditional model:
##  Groups     Name        Variance Std.Dev.
##  species_id (Intercept) 0.30082  0.5485
##  sample_id  (Intercept) 0.02217  0.1489
##  subject_id (Intercept) 0.06698  0.2588
## Number of obs: 15355, groups:  species_id, 184; sample_id, 467; subject_id, 249
##
## Overdispersion parameter for truncated_nbinom1 family (): 1.26e-05
##
## Conditional model:
##                  Estimate Std. Error z value Pr(>|z|)
## (Intercept)      -0.32627    0.05587  -5.840 5.23e-09 ***
## class_nb         -0.06721    0.01819  -3.694 0.000221 ***
## total_reads_orig  0.08639    0.02075   4.163 3.14e-05 ***
## ---
## Signif. codes:  0 '***' 0.001 '**' 0.01 '*' 0.05 '.' 0.1 ' ' 1
```

#### 9.3. Strain count in a focal species as a function of order richness

```
##  Family: truncated_nbinom2  ( log )
## Formula:
## strain_nb ~ order_nb + total_reads_orig + (order_nb | species_id) +
##     (1 | sample_id) + (1 | subject_id)
## Data: datsc2
##
##      AIC      BIC   logLik deviance df.resid
##  24922.1  24990.8 -12452.0  24904.1    15346
##
## Random effects:
##
## Conditional model:
##  Groups     Name        Variance Std.Dev. Corr
##  species_id (Intercept) 0.298733 0.5466
##             order_nb    0.001325 0.0364   0.99
##  sample_id  (Intercept) 0.021727 0.1474
##  subject_id (Intercept) 0.059083 0.2431
## Number of obs: 15355, groups:  species_id, 184; sample_id, 467; subject_id, 249
##
## Overdispersion parameter for truncated_nbinom2 family (): 2.16e+04
##
## Conditional model:
##                  Estimate Std. Error z value Pr(>|z|)
## (Intercept)      -0.32528    0.05516  -5.897 3.71e-09 ***
## order_nb         -0.11970    0.02012  -5.951 2.67e-09 ***
## total_reads_orig  0.11110    0.02127   5.224 1.75e-07 ***
## ---
## Signif. codes:  0 '***' 0.001 '**' 0.01 '*' 0.05 '.' 0.1 ' ' 1
```

#### 9.4. Strain count in a focal species as a function of family richness

```
##  Family: truncated_nbinom1  ( log )
## Formula:
## strain_nb ~ family_nb + total_reads_orig + (family_nb | species_id) +
##     (1 | sample_id) + (1 | subject_id)
## Data: datsc2
##
##      AIC      BIC   logLik deviance df.resid
##  24915.0  24983.7 -12448.5  24897.0    15346
##
## Random effects:
##
## Conditional model:
##  Groups     Name        Variance  Std.Dev. Corr
##  species_id (Intercept) 0.2987158 0.54655
##             family_nb   0.0005029 0.02242  0.89
##  sample_id  (Intercept) 0.0189422 0.13763
##  subject_id (Intercept) 0.0590925 0.24309
## Number of obs: 15355, groups:  species_id, 184; sample_id, 467; subject_id, 249
##
## Overdispersion parameter for truncated_nbinom1 family (): 1.71e-05
##
## Conditional model:
##                  Estimate Std. Error z value Pr(>|z|)
## (Intercept)      -0.31927    0.05516  -5.788 7.11e-09 ***
## family_nb        -0.12908    0.01937  -6.664 2.66e-11 ***
## total_reads_orig  0.11002    0.02038   5.398 6.73e-08 ***
## ---
## Signif. codes:  0 '***' 0.001 '**' 0.01 '*' 0.05 '.' 0.1 ' ' 1
```

#### 9.5. Strain count in a focal species as a function of genus richness

```
##  Family: truncated_nbinom1  ( log )
## Formula:
## strain_nb ~ genus_nb + total_reads_orig + (genus_nb | species_id) +
##     (1 | sample_id) + (1 | subject_id)
## Data: datsc2
##
##      AIC      BIC   logLik deviance df.resid
##  24943.1  25011.8 -12462.5  24925.1    15346
##
## Random effects:
##
## Conditional model:
##  Groups     Name        Variance  Std.Dev. Corr
##  species_id (Intercept) 0.3011560 0.54878
##             genus_nb    0.0001142 0.01069  0.86
##  sample_id  (Intercept) 0.0211474 0.14542
##  subject_id (Intercept) 0.0671252 0.25909
## Number of obs: 15355, groups:  species_id, 184; sample_id, 467; subject_id, 249
##
## Overdispersion parameter for truncated_nbinom1 family (): 1.82e-05
##
## Conditional model:
##                  Estimate Std. Error z value Pr(>|z|)
## (Intercept)      -0.32475    0.05580  -5.820 5.90e-09 ***
## genus_nb         -0.08379    0.02090  -4.009 6.09e-05 ***
## total_reads_orig  0.10105    0.02174   4.648 3.36e-06 ***
## ---
## Signif. codes:  0 '***' 0.001 '**' 0.01 '*' 0.05 '.' 0.1 ' ' 1
```

### 10. DBD over time in the gut (HMP time series)

#### 10.1. Polymorphism change in a focal species as a function of community diversity at the earlier time point

##### 10.1.1. Polymorphism change in a focal species as a function of Shannon diversity at the earlier time point

```
##
## Family: gaussian
## Link function: identity
##
## Formula:
## log.delta_pi ~ s(total_reads_orig1) + s(alpha_div_tp1) + s(species,
##     alpha_div_tp1, bs = "fs") + s(sample_tp1, bs = "re") + s(subject,
##     bs = "re")
##
## Parametric coefficients:
##               Estimate Std. Error t value Pr(>|t|)
## (Intercept) -4.943e-06  7.213e-05  -0.069    0.945
##
## Approximate significance of smooth terms:
##                                edf  Ref.df     F  p-value
## s(total_reads_orig1)     1.000e+00   1.000 1.236 0.266297
## s(alpha_div_tp1)         2.286e+00   2.757 1.365 0.376384
## s(species,alpha_div_tp1) 2.099e-04 591.000 0.000 0.990102
## s(sample_tp1)            4.552e+01 215.000 0.297 0.000338 ***
## s(subject)               8.611e-04 159.000 0.000 0.721101
## ---
## Signif. codes:  0 '***' 0.001 '**' 0.01 '*' 0.05 '.' 0.1 ' ' 1
##
## R-sq.(adj) =  0.0227   Deviance explained = 3.83%
## GCV = 1.2171e-05  Scale est. = 1.1973e-05  n = 3063
```

##### 10.1.2. Polymorphism change in a focal species as a function of species richness at the earlier time point

```
##
## Family: gaussian
## Link function: identity
##
## Formula:
## log.delta_pi ~ s(total_reads_orig1) + s(richness_tp1) + s(species,
##     richness_tp1, bs = "fs") + s(sample_tp1, bs = "re") + s(subject,
##     bs = "re")
##
## Parametric coefficients:
##               Estimate Std. Error t value Pr(>|t|)
## (Intercept) -1.072e-05  7.415e-05  -0.145    0.885
##
## Approximate significance of smooth terms:
##                               edf  Ref.df     F  p-value
## s(total_reads_orig1)    1.000e+00   1.000 1.666 0.196889
## s(richness_tp1)         1.361e+00   1.585 0.256 0.606808
## s(species,richness_tp1) 1.051e-04 583.000 0.000 0.999966
## s(sample_tp1)           5.395e+01 215.000 0.357 0.000178 ***
## s(subject)              1.119e-04 159.000 0.000 0.821816
## ---
## Signif. codes:  0 '***' 0.001 '**' 0.01 '*' 0.05 '.' 0.1 ' ' 1
##
## R-sq.(adj) =  0.025   Deviance explained =  4.3%
## GCV = 1.2172e-05  Scale est. = 1.1945e-05  n = 3063
```

##### 10.1.3. Polymorphism change in a focal species as a function of the rarefied species richness at the earlier time point

```
##
## Family: gaussian
## Link function: identity
##
## Formula:
## log.delta_pi ~ s(richness_rare_tp1) + s(total_reads_orig) + s(species,
##     richness_rare_tp1, bs = "fs") + s(sample_tp1, bs = "re") +
##     s(subject, bs = "re")
##
## Parametric coefficients:
##               Estimate Std. Error t value Pr(>|t|)
## (Intercept) -9.546e-06  7.241e-05  -0.132    0.895
##
## Approximate significance of smooth terms:
##                                    edf  Ref.df     F  p-value
## s(richness_rare_tp1)          1.000025   1.000 0.319 0.572048
## s(total_reads_orig)           1.005210   1.009 0.971 0.324110
## s(species,richness_rare_tp1)  4.108641 558.000 0.009 0.215833
## s(sample_tp1)                46.996360 215.000 0.312 0.000169 ***
## s(subject)                    0.001492 159.000 0.000 0.711795
## ---
## Signif. codes:  0 '***' 0.001 '**' 0.01 '*' 0.05 '.' 0.1 ' ' 1
##
## R-sq.(adj) =  0.0241   Deviance explained =  4.1%
## GCV = 1.2171e-05  Scale est. = 1.1956e-05  n = 3063
```

#### 10.2. Gene gain in a focal species as a function of community diversity at the earlier time point

##### 10.2.1. Gene gain in a focal species as a function of Shannon diversity at the earlier time point

```
##  Family: nbinom1  ( log )
## Formula:
## num_gene_gains ~ alpha_div_tp1 + total_reads_orig1 + (alpha_div_tp1 |
##     species) + (1 | subject) + (1 | sample_tp1)
## Data: datsc
##
##      AIC      BIC   logLik deviance df.resid
##   1516.0   1562.5   -749.0   1498.0     1287
##
## Random effects:
##
## Conditional model:
##  Groups     Name          Variance Std.Dev. Corr
##  species    (Intercept)   0.28885  0.5374
##             alpha_div_tp1 0.08704  0.2950   -0.37
##  subject    (Intercept)   2.12132  1.4565
##  sample_tp1 (Intercept)   1.00077  1.0004
## Number of obs: 1296, groups:  species, 54; subject, 154; sample_tp1, 210
##
## Overdispersion parameter for nbinom1 family ():  121
##
## Conditional model:
##                   Estimate Std. Error z value Pr(>|z|)
## (Intercept)       -0.36000    0.41245  -0.873    0.383
## alpha_div_tp1      0.07365    0.21687   0.340    0.734
## total_reads_orig1 -0.08165    0.23305  -0.350    0.726
```

##### 10.2.2. Gene gain in a focal species as a function of species richness at the earlier time point

```
##  Family: nbinom1  ( log )
## Formula:
## num_gene_gains ~ richness_tp1 + total_reads_orig1 + (1 | species) +
##     (1 | subject) + (1 | sample_tp1)
## Data: datsc
##
##      AIC      BIC   logLik deviance df.resid
##   1513.1   1549.2   -749.5   1499.1     1289
##
## Random effects:
##
## Conditional model:
##  Groups     Name        Variance Std.Dev.
##  species    (Intercept) 0.2914   0.5398
##  subject    (Intercept) 2.1331   1.4605
##  sample_tp1 (Intercept) 0.8869   0.9418
## Number of obs: 1296, groups:  species, 54; subject, 154; sample_tp1, 210
##
## Overdispersion parameter for nbinom1 family ():  123
##
## Conditional model:
##                   Estimate Std. Error z value Pr(>|z|)
## (Intercept)       -0.28956    0.39887  -0.726    0.468
## richness_tp1       0.09735    0.19348   0.503    0.615
## total_reads_orig1 -0.14904    0.24677  -0.604    0.546
```

##### 10.2.3. Gene gain in a focal species as a function of rarefied richness at the earlier time point

```
##  Family: nbinom1  ( log )
## Formula:
## num_gene_gains ~ richness_rare_tp1 + (1 | species) + (1 | subject) +
##     (1 | sample_tp1)
## Data: datsc
##
##      AIC      BIC   logLik deviance df.resid
##   1511.1   1542.1   -749.5   1499.1     1290
##
## Random effects:
##
## Conditional model:
##  Groups     Name        Variance Std.Dev.
##  species    (Intercept) 0.2925   0.5408
##  subject    (Intercept) 2.1466   1.4651
##  sample_tp1 (Intercept) 0.9015   0.9495
## Number of obs: 1296, groups:  species, 54; subject, 154; sample_tp1, 210
##
## Overdispersion parameter for nbinom1 family ():  123
##
## Conditional model:
##                   Estimate Std. Error z value Pr(>|z|)
## (Intercept)        -0.2833     0.3971  -0.713    0.476
## richness_rare_tp1   0.1127     0.1691   0.666    0.505
```

#### 10.3. Gene loss in a focal species as a function of community diversity at the earlier time point

##### 10.3.1. Gene loss in a focal species as a function of Shannon diversity at the earlier time point

```
##  Family: nbinom2  ( log )
## Formula:
## num_gene_losses ~ alpha_div_tp1 + total_reads_orig1 + (alpha_div_tp1 |
##     species) + (1 | subject) + (1 | sample_tp1)
## Data: datsc
##
##      AIC      BIC   logLik deviance df.resid
##   1696.2   1742.7   -839.1   1678.2     1287
##
## Random effects:
##
## Conditional model:
##  Groups     Name          Variance Std.Dev. Corr
##  species    (Intercept)    1.7511  1.3233
##             alpha_div_tp1  0.2719  0.5214   -0.57
##  subject    (Intercept)    2.3094  1.5197
##  sample_tp1 (Intercept)   10.3411  3.2158
## Number of obs: 1296, groups:  species, 54; subject, 154; sample_tp1, 210
##
## Overdispersion parameter for nbinom2 family (): 0.111
##
## Conditional model:
##                   Estimate Std. Error z value Pr(>|z|)
## (Intercept)        -3.5455     0.8326  -4.259 2.06e-05 ***
## alpha_div_tp1       0.8360     0.3950   2.117   0.0343 *
## total_reads_orig1   0.8956     0.5299   1.690   0.0910 .
## ---
## Signif. codes:  0 '***' 0.001 '**' 0.01 '*' 0.05 '.' 0.1 ' ' 1
```

##### 10.3.2. Gene loss in a focal species as a function of species richness at the earlier time point

```
##  Family: nbinom2  ( log )
## Formula:
## num_gene_losses ~ richness_tp1 + total_reads_orig1 + (1 | species) +
##     (1 | subject) + (1 | sample_tp1)
## Data: datsc
##
##      AIC      BIC   logLik deviance df.resid
##   1695.1   1731.2   -840.5   1681.1     1289
##
## Random effects:
##
## Conditional model:
##  Groups     Name        Variance Std.Dev.
##  species    (Intercept) 1.577    1.256
##  subject    (Intercept) 3.432    1.852
##  sample_tp1 (Intercept) 8.125    2.850
## Number of obs: 1296, groups:  species, 54; subject, 154; sample_tp1, 210
##
## Overdispersion parameter for nbinom2 family (): 0.102
##
## Conditional model:
##                   Estimate Std. Error z value Pr(>|z|)
## (Intercept)        -3.3062     0.7822  -4.227 2.37e-05 ***
## richness_tp1        0.7851     0.3751   2.093   0.0364 *
## total_reads_orig1   0.3357     0.5010   0.670   0.5028
## ---
## Signif. codes:  0 '***' 0.001 '**' 0.01 '*' 0.05 '.' 0.1 ' ' 1
```

##### 10.3.3. Gene loss in a focal species as a function of rarefied richness at the earlier time point

```
##  Family: nbinom1  ( log )
## Formula:
## num_gene_losses ~ richness_rare_tp1 + total_reads_orig1 + (richness_rare_tp1 |
##     species) + (1 | subject) + (1 | sample_tp1)
## Data: datsc
##
##      AIC      BIC   logLik deviance df.resid
##   1709.6   1756.1   -845.8   1691.6     1287
##
## Random effects:
##
## Conditional model:
##  Groups     Name              Variance Std.Dev. Corr
##  species    (Intercept)       0.22747  0.4769
##             richness_rare_tp1 0.09539  0.3088   -0.58
##  subject    (Intercept)       0.94132  0.9702
##  sample_tp1 (Intercept)       1.01547  1.0077
## Number of obs: 1296, groups:  species, 54; subject, 154; sample_tp1, 210
##
## Overdispersion parameter for nbinom1 family ():  175
##
## Conditional model:
##                   Estimate Std. Error z value Pr(>|z|)
## (Intercept)         0.4533     0.3213   1.411   0.1583
## richness_rare_tp1   0.3501     0.1708   2.050   0.0404 *
## total_reads_orig1   0.2109     0.1827   1.155   0.2483
## ---
## Signif. codes:  0 '***' 0.001 '**' 0.01 '*' 0.05 '.' 0.1 ' ' 1
```

### 11. DBD over time in the gut (Poyet time series)

#### 11.1 polymorphism rate variation as a function of Shannon diversity at the earlier time point

```
##
## Family: gaussian
## Link function: identity
##
## Formula:
## log.delta_pi ~ ti(alpha_div_tp1, lag) + s(read_count) + s(species,
##     bs = "re") + s(subject, bs = "re") + s(sample_tp1, bs = "re") +
##     s(sample_tp2, bs = "re")
##
## Parametric coefficients:
##              Estimate Std. Error t value Pr(>|t|)
## (Intercept) 0.0003851  0.0006152   0.626    0.531
##
## Approximate significance of smooth terms:
##                           edf  Ref.df         F  p-value
## ti(alpha_div_tp1,lag)   5.402   6.915     2.325   0.0234 *
## s(read_count)           1.006   1.006     1.633   0.1998
## s(species)             13.552  14.000  3179.569 2.83e-07 ***
## s(subject)              1.897   3.000 93229.029   0.2388
## s(sample_tp1)         352.188 362.000   364.322 3.55e-16 ***
## s(sample_tp2)         350.584 359.000  2950.333 5.61e-15 ***
## ---
## Signif. codes:  0 '***' 0.001 '**' 0.01 '*' 0.05 '.' 0.1 ' ' 1
##
## R-sq.(adj) =  0.655   Deviance explained = 66.3%
## -ML = -1.6937e+05  Scale est. = 1.3729e-06  n = 32113
```

#### 11.2 polymorphism rate variation as a function of rarefied richness at the earlier time point

```
##
## Family: gaussian
## Link function: identity
##
## Formula:
## log.delta_pi ~ ti(richness_rare_tp1, lag) + s(read_count) + s(species,
##     bs = "re") + s(subject, bs = "re") + s(sample_tp1, bs = "re") +
##     s(sample_tp2, bs = "re")
##
## Parametric coefficients:
##              Estimate Std. Error t value Pr(>|t|)
## (Intercept) 0.0004998  0.0005718   0.874    0.382
##
## Approximate significance of smooth terms:
##                                 edf Ref.df        F  p-value
## ti(richness_rare_tp1,lag)   0.01718     16    0.001    0.689
## s(read_count)               0.31775      9 4909.967    0.225
## s(species)                 13.54785     14 2995.679 6.36e-10 ***
## s(subject)                  0.02014      3   28.740    0.290
## s(sample_tp1)             353.84479    363 1114.032 1.31e-06 ***
## s(sample_tp2)             351.65462    359 6153.476 3.90e-07 ***
## ---
## Signif. codes:  0 '***' 0.001 '**' 0.01 '*' 0.05 '.' 0.1 ' ' 1
##
## R-sq.(adj) =  0.655   Deviance explained = 66.3%
## -ML = -1.6936e+05  Scale est. = 1.3735e-06  n = 32113
```

#### 11.3 Gene gain in a focal species as a function of Shannon diversity at the earlier time point

```
##  Family: nbinom2  ( log )
## Formula:
## num_gene_gains ~ alpha_div_tp1 * lag + read_count + (1 | species) +
##     (1 | subject) + (1 | sample_tp1) + (1 | sample_tp2)
## Data: datsc
##
##      AIC      BIC   logLik deviance df.resid
##  13058.3  13142.0  -6519.1  13038.3    32103
##
## Random effects:
##
## Conditional model:
##  Groups     Name        Variance Std.Dev.
##  species    (Intercept) 4.8180   2.195
##  subject    (Intercept) 3.8308   1.957
##  sample_tp1 (Intercept) 0.5373   0.733
##  sample_tp2 (Intercept) 6.3800   2.526
## Number of obs: 32113, groups:
## species, 15; subject, 4; sample_tp1, 364; sample_tp2, 360
##
## Overdispersion parameter for nbinom2 family (): 2.54
##
## Conditional model:
##                   Estimate Std. Error z value Pr(>|z|)
## (Intercept)       -5.86584    1.32874  -4.415 1.01e-05 ***
## alpha_div_tp1     -0.50462    0.07201  -7.008 2.42e-12 ***
## lag                0.64204    0.06529   9.834  < 2e-16 ***
## read_count        -0.06774    0.05990  -1.131    0.258
## alpha_div_tp1:lag -0.21991    0.03632  -6.054 1.41e-09 ***
## ---
## Signif. codes:  0 '***' 0.001 '**' 0.01 '*' 0.05 '.' 0.1 ' ' 1
```

#### 11.4 Gene gain in a focal species as a function of rarefied richness at the earlier time point

```
##  Family: nbinom2  ( log )
## Formula:          num_gene_gains ~ richness_rare_tp1 * lag + read_count + (1 |
##     species) + (1 | subject) + (1 | sample_tp1) + (1 | sample_tp2)
## Data: datsc1
##
##      AIC      BIC   logLik deviance df.resid
##  13120.8  13204.5  -6550.4  13100.8    32103
##
## Random effects:
##
## Conditional model:
##  Groups     Name        Variance Std.Dev.
##  species    (Intercept) 5.1570   2.2709
##  subject    (Intercept) 3.6070   1.8992
##  sample_tp1 (Intercept) 0.6188   0.7866
##  sample_tp2 (Intercept) 6.2895   2.5079
## Number of obs: 32113, groups:
## species, 15; subject, 4; sample_tp1, 364; sample_tp2, 360
##
## Overdispersion parameter for nbinom2 family (): 2.41
##
## Conditional model:
##                       Estimate Std. Error z value Pr(>|z|)
## (Intercept)           -6.13441    1.32483  -4.630 3.65e-06 ***
## richness_rare_tp1     -0.05682    0.06514  -0.872 0.383100
## lag                    0.71254    0.06831  10.431  < 2e-16 ***
## read_count            -0.17279    0.06142  -2.813 0.004908 **
## richness_rare_tp1:lag -0.13105    0.03661  -3.580 0.000343 ***
## ---
## Signif. codes:  0 '***' 0.001 '**' 0.01 '*' 0.05 '.' 0.1 ' ' 1
```

#### 11.5 Gene loss in a focal species as a function of Shannon diversity at the earlier time point

```
##  Family: nbinom2  ( log )
## Formula:
## num_gene_losses ~ alpha_div_tp1 * lag + read_count + (1 | species) +
##     (1 | subject) + (1 | sample_tp1) + (1 | sample_tp2)
## Data: datsc
##
##      AIC      BIC   logLik deviance df.resid
##  13168.6  13252.4  -6574.3  13148.6    32103
##
## Random effects:
##
## Conditional model:
##  Groups     Name        Variance Std.Dev.
##  species    (Intercept) 1.7682   1.3297
##  subject    (Intercept) 3.0006   1.7322
##  sample_tp1 (Intercept) 4.6178   2.1489
##  sample_tp2 (Intercept) 0.3656   0.6047
## Number of obs: 32113, groups:
## species, 15; subject, 4; sample_tp1, 364; sample_tp2, 360
##
## Overdispersion parameter for nbinom2 family ():  3.8
##
## Conditional model:
##                   Estimate Std. Error z value Pr(>|z|)
## (Intercept)       -6.32440    1.05926  -5.971 2.36e-09 ***
## alpha_div_tp1      0.81713    0.16397   4.984 6.24e-07 ***
## lag                0.22256    0.04946   4.500 6.79e-06 ***
## read_count        -0.07615    0.15416  -0.494  0.62132
## alpha_div_tp1:lag  0.10992    0.04017   2.736  0.00621 **
## ---
## Signif. codes:  0 '***' 0.001 '**' 0.01 '*' 0.05 '.' 0.1 ' ' 1
```

#### 11.6 Gene loss in a focal species as a function of rarefied richness at the earlier time point

```
##  Family: nbinom2  ( log )
## Formula:
## num_gene_losses ~ richness_rare_tp1 * lag + read_count + (1 |
##     species) + (1 | subject) + (1 | sample_tp1) + (1 | sample_tp2)
## Data: datsc1
##
##      AIC      BIC   logLik deviance df.resid
##  13188.7  13272.4  -6584.3  13168.7    32103
##
## Random effects:
##
## Conditional model:
##  Groups     Name        Variance Std.Dev.
##  species    (Intercept) 1.6549   1.2864
##  subject    (Intercept) 3.3111   1.8196
##  sample_tp1 (Intercept) 4.9436   2.2234
##  sample_tp2 (Intercept) 0.3674   0.6061
## Number of obs: 32113, groups:
## species, 15; subject, 4; sample_tp1, 364; sample_tp2, 360
##
## Overdispersion parameter for nbinom2 family ():  3.8
##
## Conditional model:
##                       Estimate Std. Error z value Pr(>|z|)
## (Intercept)           -5.89552    1.07721  -5.473 4.43e-08 ***
## richness_rare_tp1      0.49358    0.16024   3.080  0.00207 **
## lag                    0.21302    0.05107   4.171 3.03e-05 ***
## read_count             0.04467    0.15553   0.287  0.77395
## richness_rare_tp1:lag  0.04897    0.04520   1.083  0.27860
## ---
## Signif. codes:  0 '***' 0.001 '**' 0.01 '*' 0.05 '.' 0.1 ' ' 1
```
