## Supplementary File 1 for "Community diversity is associated with intra-species genetic diversity and gene loss in the human gut microbiome"

**Supplementary tables**

**Table a.** **Generalized additive models of polymorphism rate (at synonymous sites) in a focal species as a function of community diversity**. We report the response, the predictors (fixed effects), the corresponding *P*-value (Chi-square test), *P-*value corresponding to species random effect on the slope and the adjusted R^2^ of the model. RE: random effect on the slope. The R syntax and further details about these models are reported in Supplementary File 2 section 1.

| **response** | **predictor** | ***P*** | ***P* (species RE)** | **R^2^** |
| --- | --- | --- | --- | --- |
| Polymorphism rate | Shannon diversity | 0.031 | < 2e-16 | 0.198 |
|  | coverage | 8.83e-04 |  |  |
| Polymorphism rate | Species richness (all) | 0.017 | < 2e-16 | 0.191 |
|  | coverage | 2.32e-06 |  |  |
| Polymorphism rate | Species richness (raref)  coverage | 2.63e-04  0.02 | < 2e-16 | 0.19 |

**Table b.** **Generalized additive models of polymorphism rate (at non-synonymous sites) in a focal species as a function of community diversity**. We report the response, the predictors (fixed effects), the corresponding *P*-value (Chi-square test), *P*-value corresponding to species random effect on the slope and the adjusted R^2^ of the model. RE: random effect on the slope. The R syntax and further details about these models are reported in Supplementary File 2 section 4.

| **response** | **predictor** | ***P*** | ***P* (species RE)** | **R^2^** |
| --- | --- | --- | --- | --- |
| Polymorphism rate | Shannon diversity | 0.882 | < 2e-16 | 0.248 |
|  | coverage | < 2e-16 |  |  |
| Polymorphism rate | Species richness (all) | 0.093 | < 2e-16 | 0.243 |
|  | coverage | < 2e-16 |  |  |
| Polymorphism rate | Species richness (raref)  coverage | 0.26  < 2e-16 | < 2e-16 | 0.244 |

**Table c.** **Generalized additive models of polymorphism rate (at synonymous sites) in a focal species as a function of community diversity at higher taxonomic levels**. We report the taxonomic level, the predictors (fixed effects), the corresponding *P*-value (Chi-square test), the *P*-value corresponding to species random effect on the slope (Chi-square test) and the adjusted R^2^ of the model. RE: random effect on the slope. The R syntax and further model details models are reported in Supplementary File 2 sections 2-3.

| **Taxonomic level** | **Predictor** | ***P*** | ***P* (species RE)** | **R^2^** |
| --- | --- | --- | --- | --- |
| phyla | Shannon div | 0.528 | < 2e-16 | 0.189 |
|  | coverage | 2.24e-04 |  |  |
| class | Shannon div | 0.518 | < 2e-16 | 0.189 |
|  | coverage | 2.27e-04 |  |  |
| order | Shannon div | 0.395 | < 2e-16 | 0.189 |
|  | coverage | 2.35e-04 |  |  |
| family | Shannon div | 0.031 | < 2e-16 | 0.192 |
|  | coverage | 8.07e-04 |  |  |
| genus | Shannon div | 0.011 | < 2e-16 | 0.193 |
|  | coverage | 8.6e-04 |  |  |
| phyla | Richness | 0.006 | < 2e-16 | 0.196 |
|  | coverage | 2.07e-05 |  |  |
| class | Richness | 0.011 | < 2e-16 | 0.194 |
|  | coverage | 1.23e-05 |  |  |
| order | Richness | 0.018 | < 2e-16 | 0.19 |
|  | coverage | 1.01e-05 |  |  |
| family | Richness | 0.002 | < 2e-16 | 0.191 |
|  | coverage | 3.37e-06 |  |  |
| genus | Richness | 0.004 | < 2e-16 | 0.193 |
|  | coverage | 2.42e-06 |  |  |

**Table d.** **Generalized additive models of polymorphism rate (at non-synonymous sites) in a focal species as a function of community diversity at higher taxonomic levels**. We report the taxonomic level, the predictors (fixed effects), the corresponding *P*-value (Chi-square test), the *P*-value corresponding to species random effect on the slope (Chi-square test) and the adjusted R^2^ of the model. RE: random effect on the slope. The R syntax and further model details are reported in the Supplementary File 2 sections 5-6.

| **Taxonomic level** | **Predictor** | ***P*** | ***P* (species RE)** | **R^2^** |
| --- | --- | --- | --- | --- |
| phyla | Shannon div | 0.284 | < 2e-16 | 0.241 |
|  | coverage | < 2e-16 |  |  |
| class | Shannon div | 0.286 | < 2e-16 | 0.24 |
|  | coverage | < 2e-16 |  |  |
| order | Shannon div | 0.138 | < 2e-16 | 0.242 |
|  | coverage | < 2e-16 |  |  |
| family | Shannon div | 0.842 | < 2e-16 | 0.244 |
|  | coverage | < 2e-16 |  |  |
| genus | Shannon div | 0.871 | < 2e-16 | 0.247 |
|  | coverage | < 2e-16 |  |  |
| phyla | Richness | 0.0003 | < 2e-16 | 0.246 |
|  | coverage | < 2e-16 |  |  |
| class | Richness | 0.017 | < 2e-16 | 0.244 |
|  | coverage | < 2e-16 |  |  |
| order | Richness | 6.11e-04 | < 2e-16 | 0.244 |
|  | coverage | < 2e-16 |  |  |
| family | Richness | 0.241 | < 2e-16 | 0.243 |
|  | coverage | < 2e-16 |  |  |
| genus | Richness | 0.122 | < 2e-16 | 0.243 |
|  | coverage | < 2e-16 |  |  |

**Table e**. **Results from Generalized Linear Mixed Models with community diversity as the predictor of strain count in a focal species**. Each row reports the response, the predictors (fixed effects), the effect and its corresponding *P*-value (LRT, drop1 function from the R package stats), the standard deviation on the slope for species random effect, and the difference in Akaike information criterion (dAIC) between the models and null models (exactly the same model but with no fixed terms) as well as the corresponding *P*-value (LRT, anova function from stats package in R). The adjusted r-squared is not reported for these models because truncated negative binomial distributions are not yet implemented in the r2 R function (performance package). SD: among species standard deviation on the slope. The R syntax and further details about these models are reported in the Supplementary File 2 section 7.

| **Response** | **Predictor** | **Effect** | **P (LRT)** | **SD Species (slope)** | **LRT/null model** | |
| --- | --- | --- | --- | --- | --- | --- |
|  |  |  |  |  | **dAIC** | ***P*** |
| strain count | Shannon diversity | 0.08 | 1.549e-04 | 0.068 | 26 | 2.6e-07 |
|  | coverage | 0.084 | 2.876e-05 |  |  |  |
| strain count | Species richness (all) | -0.089 | 1.500e-05 | 0.024 | 30 | 3.727e-08 |
|  | coverage | 0.118 | 6.37e-08 |  |  |  |
| strain count | Species richness (rarefied) | -0.041 | 0.037 | 0.009 | 2 | 0.037 |

**Table f.** **Generalized Linear Mixed Models with community diversity at higher taxonomic levels as the predictor of strain count in a focal species.** Each row reports the taxonomic level, the predictors (fixed effects), the effect and its corresponding *P*-value (likelihood-ratio test, drop1 function from the stats R package), among species standard deviation on the slope (SD) and the difference in Akaike information criterion between the full and null models (exactly the same model but with no fixed terms) and the corresponding *P*-value (LRT, anova function from stats package in R). The adjusted r-squared is not reported for these models because truncated negative binomial distributions are not yet implemented in the r2 R function (performance package). ns=not significant (LRT, anova function from the stats R package). The R syntax and further details about these models are reported in the Supplementary File 2 sections 8-9.

| **Taxonomic level** | **Predictor** | **Effect** | **P(LRT)** | **SD species (slope)** | **LRT/null model** | | |
| --- | --- | --- | --- | --- | --- | --- | --- |
|  |  |  |  |  | **dAIC** | | ***P*** |
| phyla | Shannon | 0.03 | 0.123 | 0.04 | Null model-Conv problem | | |
|  | coverage | 0.065 | 0.001 |  |  | | |
| class | Shannon | 0.032 | 0.103 | 0.043 | 8 | 0.002 | |
|  | coverage | 0.066 | 0.001 |  |  |  | |
| order | Shannon | 0.034 | 0.008 | 0.045 | 9 | 0.002 | |
|  | coverage | 0.028 | 0.01 |  |  |  | |
| family | Shannon | 0.03 | 0.129 | 0.036 | 8 | 0.002 | |
|  | coverage | 0.066 | 0.001 |  |  |  | |
| genus | Shannon | 0.057 | 0.003 | 0.057 | 14 | 1.49e-04 | |
|  | coverage | 0.065 | 0.002 |  |  |  | |
| phyla | richness | -0.048 | 2.204e-05 | ns | 21 | 4.722e-06 | |
|  | coverage | 0.036 | 0.001 |  |  |  | |
| class | richness | -0.067 | 0.0002 | ns | 20 | 6.124e-06 | |
|  | coverage | 0.086 | 3.14e-05 |  |  |  | |
| order | richnes | -0.12 | 3.318e-09 | 0.0364 | the null model did not converge | | |
|  | coverage | 0.111 | 2.538e-07 |  |  | | |
| family | richness | -0.129 | 3.941e-11 | 0.022 | 51 | 1.453e-12 | |
|  | coverage | 0.11 | 9.081e-08 |  |  |  | |
| genus | richness | -0.084 | 6.09e-05 | 0.01 | 23 | 1.67e-06 | |
|  | coverage | 0.101 | 3.36e-06 |  |  |  | |

**Table g.** **GLMMs predicting gene loss between two HMP time points as a function of community diversity at the earlier time point**. Each row reports the response, the predictors (fixed effects), the effect and its corresponding *P*-value (likelihood-ratio test, drop1 function from the stats R package), among species standard deviation on the slope (SD), and the difference in Akaike information criterion between the full and null models (exactly the same model but with no fixed terms) and the corresponding and *P*-value (LRT, anova function from stats package in R). R^2^ is the adjusted R-squared of the model (r2 function from the R package performance) (Marg: marginal R^2^=measure of the variance explained only by fixed effects, cond: conditional R^2^ = measure of the variance explained by the entire model). SD: among species standard deviation on the slope. The R syntax and further details about these models are reported in the Supplementary File 2 section 10.3.

| **Response** | **Predictor** | **Effect** | **P (LRT)** | **SD Species (slope)** | **LRT/null model** | | **R^2^** | |
| --- | --- | --- | --- | --- | --- | --- | --- | --- |
|  |  |  |  |  | **dAIC** | ***P*** | **Marg** | **Cond** |
| Gene loss | Shannon tp1 | 0.836 | 0.028 | 0.5214 | 4 | 0.02 | 0.069 | 0.821 |
|  | Coverage tp1 | 0.896 | 0.055 |  |  |  |  |  |
| Gene loss | Richness tp1 (all) | 0.785 | 0.034 | ns | 3.1 | 0.028 | 0.055 | 0.805 |
|  | Coverage tp1 | 0.336 | 0.487 |  |  |  |  |  |
| Gene loss | Richness tp1 (rarefied) | 0.35 | 0.049 | ns | 0.7 | 0.094 | 0.019 | 0.335 |
|  | Coverage tp1 | 0.211 | 0.243 |  |  |  |  |  |

**Table h. GAM predicting polymorphism change as a function of the initial diversity in Poyet time series**. We report the response, the predictors, the related *P*-value (Chi-square test) and the adjusted R^2^ of the model. The GAMs were fitted with Gaussian error distribution and log transformed polymorphism change as the response, with Shannon index and rarefied richness as the predictors. The R syntax and further details about the model are reported in Supplementary File 2 sections 11.1-11.2.

| **Response** | **Predictor** | ***P*** | **R^2^** |
| --- | --- | --- | --- |
| Log10 1+delta pi | Shannon tp1*time lag | 0.0234 | 0.655 |
|  | Coverage tp1 | 0.1998 |  |
| Log10 1+delta pi | richness tp1*time lag | 0.689 | 0.655 |
|  | Coverage tp1 | 0.225 |  |

**Table i.** **GLMMs predicting gene loss/gain between two time points as a function of diversity at the earlier time point in Poyet time series data**. Each row reports the response, the predictors (fixed effects), the effect and its corresponding *P*-value (likelihood-ratio test, drop1 function from the stats R package. The difference in Akaike information criterion and *P*-value (LRT, anova function from stats package in R) between the full and null models (exactly the same model but with no fixed terms). R^2^ is the adjusted R-squared of the model (r2 function from performance R package) (Marg: marginal R^2^= measure of the variance explained only by fixed effects, cond: conditional R^2^ =measure of the variance explained by the entire model). The R syntax and further model details are reported in Supplementary File 2 sections 11.3-11.6.

| **Response** | **Predictor** | **Effect** | **P (LRT)** | **LRT/null model** | | **R^2^** | |
| --- | --- | --- | --- | --- | --- | --- | --- |
|  |  |  |  | **dAIC** | ***P*** | **Marg** | **Cond** |
| Gene loss | Shannon tp1*time lag | 0.11 | 0.006 | 44 | 9.54e-11 | 0.037 | 0.638 |
|  | Coverage tp1 | -0.076 | 0.621 |  |  |  |  |
| Gene loss | richness tp1*time lag | 0.005 | 0.278 | 27 | 1.49e-07 | 0.018 | 0.643 |
|  | Coverage tp1 | 0.045 | 0.774 |  |  |  |  |
| Gene gain | Shannon tp1*time lag | -0.22 | 1.11e-09 | 634 | 2.2e-16 | 0.037 | 0.713 |
|  | Coverage tp1 | 0.068 | 0.259 |  |  |  |  |
| Gene gain | richness tp1*time lag | -0.131 | 3.413e-04 | 113 | 2.2e-16 | 0.025 | 0.711 |
|  | Coverage tp1 | -0.173 | 5.247e-03 |  |  |  |  |
